# Quantifying the impact of experimental hut design on intervention evaluation outcomes and predicted reductions in vectorial capacity

**DOI:** 10.64898/2026.08.24.746696

**Authors:** Emma L Fairbanks, Alphonce Assenga, Olukayode G Odufuwa, Raphael N’Guessan, Jason Moore, Sarah J Moore

## Abstract

Experimental hut trials (EHTs) are WHO-recommended for the entomological evaluation of insecticide-treated nets (ITNs), but several hut designs are in operational use, and structural differences between them may confound efficacy predictions and limit cross-site comparability. We developed a Bayesian hierarchical framework comprising a host-seeking model, which jointly estimates biting deterrence and preprandial mortality while accounting for night-to-night variation and overdispersion, and a postprandial mortality model, which expresses hut and net effects as hazard ratios through a complementary log-log link. We applied it to a comparative trial of four hut designs (East African, West African, Ifakara and Rapley) conducted at a single site in Tanzania, evaluating eight ITNs when new and after twenty washes. Posterior estimates parameterise a vectorial capacity framework to predict reductions in transmission potential. Hut design influenced baseline mosquito behaviour and all three modes of action. Relative to the Rapley reference, baseline feeding rates were substantially lower in the East African and West African huts and closer to Rapley in the Ifakara hut. Preprandial mortality was amplified in the Ifakara hut. Comparing to previous analysis provides evidence that combining mortality before and after feeding into a single endpoint does not reliably reflect impact, supporting the decomposition of entomological outcomes into separate modes of action. Expressing modes of action as mechanism-specific parameters allows the estimates to be carried directly into transmission models. For every net, the predicted reduction in vectorial capacity was greatest in the Ifakara hut and smallest in the West African and Rapley huts. The effect of hut design on predicted impact exceeded that of washing the nets twenty times. Results indicate that the hut design under which trial data were collected should be considered when forecasting population-level effect.

## 1 Introduction

As the World Health Organisation (WHO)-recommended standard for entomological evaluation of insecticidal vector control interventions, experimental hut trials (EHTs) offer ecologically realistic conditions in which mosquitoes encounter treated nets during natural host-seeking behaviour [1, 2]. Although hut construction follows standardised principles, controlling for internal volume, entry and exit geometry and the absence of furniture, multiple distinct designs are in operational use across sub-Saharan Africa, varying in size and the configuration of mosquito entry and exit points [2].

Such structural heterogeneity has the potential to alter mosquito behaviour within huts and thereby confound predictions of efficacy (the capacity of a vector control product to reduce mosquito transmission potential), with implications for the cross-site comparability and policy utility of trial results [2–4]. Interpreting trial data is further complicated by the concurrent use of distinct hut designs across regions that differ in vector species composition and resistance profiles, making it difficult to attribute variation in outcomes to biological differences rather than methodological ones. To address this gap, Assenga et al. [3] conducted a comparative evaluation of four experimental hut designs (East African, West African, Ifakara, and Rapley) at a single site in Tanzania.

Translating entomological measurements into predictions of intervention impacts on vectorial capacity (a measure of a vector population’s ability to transmit a pathogen) requires quantitative frameworks that consider the biological structure of the underlying processes, decomposing intervention effects into distinct modes of action: biting deterrence, the prevention of feeding; preprandial mortality, death before blood-feeding; and postprandial mortality, death after blood-feeding [5]. While many models for vectorial capacity exist [6–8], those incorporating the effects of insecticide-treated nets (ITNs) typically represent net-induced mortality as a single mortality term [9–11], and models which consider the impact of all of these modes of action individually are less common [12, 13]. This distinction matters because different modes of action contribute unequally to vectorial capacity reduction [14], and systematic differences in hut architecture may lead each design to capture them to differing degrees, potentially yielding divergent estimates of population-level impact.

To address this gap, we developed a Bayesian hierarchical framework comprising two complementary models and applied it to the comparative trial of four hut designs reported by Assenga et al. [3]. The first is a host-seeking model that jointly estimates the impact on deterrence and preprandial mortality. The model explicitly accounts for day-to-day variation in mosquito activity and for differences in mosquito behaviour between hut designs, and uses a negative binomial distribution to accommodate overdispersion in the count data. The second is a postprandial mortality model. It uses a complementary log-log link, so that hut and intervention effects are expressed as hazard ratios and can be interpreted on a multiplicative scale independently of the day-specific baseline mortality. Estimates from both models are embedded in a vectorial capacity framework [12] to translate the entomological effects of each net into predicted reductions in transmission potential across a range of intervention coverage levels. This allows us to assess the extent to which experimental hut design affects the interpretation and comparability of ITN trial outcomes.

## 2 Methods

### 2.1 Data

Full details of the study design and data collection are described in Assenga et al. [3]. The study was conducted in Lupiro Village (8.385°S, 36.670°E) in the Kilombero river valley, Tanzania, where *Anopheles gambiae s*.*l*. (>99.9% *Anopheles arabiensis*) are present year-round. The local *An. arabiensis* population is resistant to pyrethroids but susceptible to chlorfenapyr.

Nine net arms were evaluated: eight ITNs (MAGNet™ ITN, MAGNet™ 2.0, Interceptor^®^ G2, Olyset^®^ Plus, PRONet^®^ Duo and three anonymised nets) alongside an untreated net negative control. Each ITN was tested in both unwashed and washed (20 standardised washes) conditions, giving 18 net arms in total. A total of 18 huts were therefore required, comprising four East African, four West African, four Rapley, and six Ifakara huts. The trial employed a 9 × 9 Latin square design, mirrored across unwashed and washed conditions, so that each net type appeared in each hut position across rounds, with unwashed and washed versions of the same net kept in the same hut type on any given night. This balanced net arms across hut designs and prevented net effects from being confounded with hut design. Within a nine-night round each net arm remained in a fixed hut, with replicate nets of the same type rotated through it nightly. Volunteers were rotated nightly to minimise bias from inter-individual differences in attractiveness. Fourteen experimental rounds of 9 nights each were completed (N=126 nights).

Each morning, mosquitoes were collected from inside the hut and exit traps. Collected mosquitoes were classified as fed and alive (FA), fed and dead (FD), unfed and dead (UD), or unfed and alive (UA). Live mosquitoes were held for up to 120 hours in a temperature-controlled room with access to 10% sugar solution to assess delayed mortality.

We use 24-hour mortality in these analyses, as mortality within this window determines whether a mosquito survives to host-seek on the following night, and because a single assessment window maintains comparability across net types.

### 2.2 Host-seeking model

The number of mosquitoes entering each hut varies considerably between nights due to factors outside experimental control, such as ambient temperature, relative humidity and local population fluctuations. We therefore model nightly counts using a negative binomial distribution, which accommodates the resulting overdispersion and high frequency of zero counts.

#### 2.2.1 Model development

Let 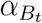 denote the baseline feeding rate on experimental day *t* and 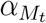 the natural preprandial mortality rate on day *t*, both on the natural scale. Both rates vary from night to night and are estimated as independent log-scale day-specific parameters,

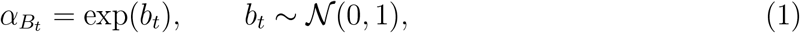

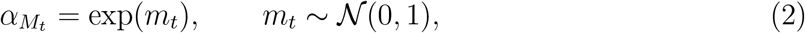

where *b*_*t*_ and *m*_*t*_ are estimated separately for each day with no assumed temporal correlation. This reflects the fact that baseline mosquito activity and natural preprandial mortality are driven by environmental conditions that do not follow a systematic trend across the experiment.

Experimental hut types differ in construction and may systematically affect mosquito behaviour independently of any net effect. Rate parameters are therefore modified by a hut-type-specific multiplicative offset on the log scale. Let *h* index hut type, with the reference hut assigned an offset of zero. The hut-adjusted baseline rates are

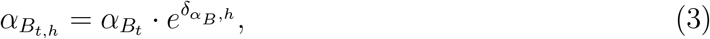

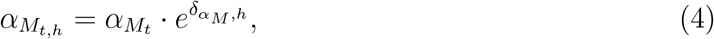

where 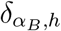 and 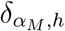 are log-scale hut offsets for baseline feeding and natural preprandial mortality, respectively. The rate of mosquitoes that either feed or die preprandially in the control arm is then

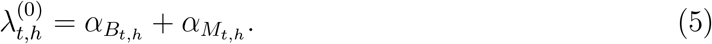

In the intervention arm, net type *n* modifies the hut-adjusted baseline rates through two net-specific effects. Deterrence scales the baseline feeding rate by a relative feeding rate 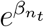, where values below one indicate reduced feeding relative to the untreated hut. Net-induced preprandial mortality adds a term 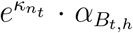 to the preprandial mortality rate; the proportionality to 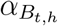 reflects that this rate is expressed relative to the number of mosquitoes that would have fed in the absence of an intervention. Log-scale hut-type offsets *δ*_*β,h*_ and *δ*_*κ,h*_ capture systematic differences between hut designs in how they modulate the relative feeding rate and preprandial mortality. The intervention rates are therefore

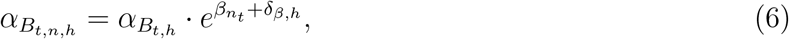

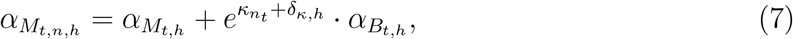

and the total rate of mosquitoes that either feed or die preprandially in the intervention arm is

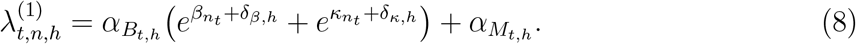

Day-to-day variation in 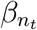 and 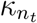 are both modelled hierarchically:

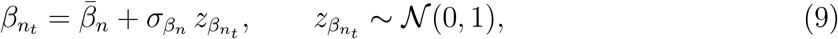

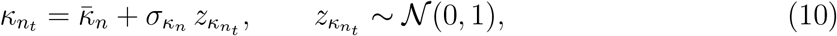

where 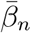 and 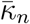 are net-type mean log-scale effects and 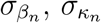 are within-net day-to-day standard deviations.

#### 2.2.2 Likelihood

We model two quantities per arm: the total count of mosquitoes that feed or die preprandially, and the conditional split between these two outcome categories.

##### Control arm

For observation *i* on day *t* in hut *h*,

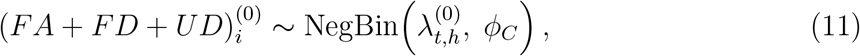

where *ϕ*_*C*_ is a shared overdispersion parameter. The negative binomial alone cannot separately identify 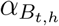 and 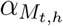 from the total count. We therefore additionally model the number of fed mosquitoes conditional on the total caught as a binomial draw,

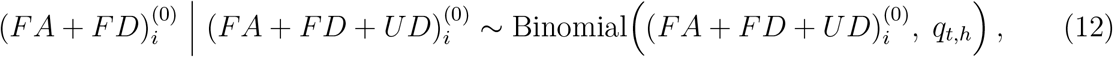

with feeding probability

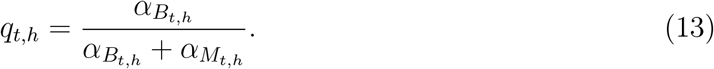

The binomial component contributes to the likelihood only when at least one mosquito is observed. Together, equations (11) and (12) identify the absolute scale of 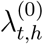 and the relative split between 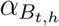 and 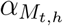 separately.

##### Intervention arm

For observation *j* with net type *n* on day *t* in hut *h*,

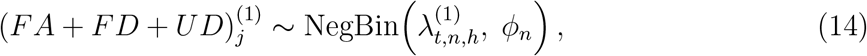

where *ϕ*_*n*_ is a net-type-specific overdispersion parameter. The number of fed mosquitoes conditional on the total is

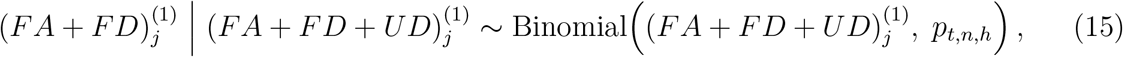

with feeding probability

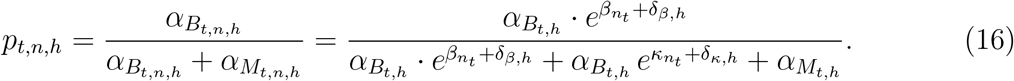

Natural preprandial mortality 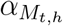 estimated from the control arm enters the intervention binomial denominator, ensuring that net-induced preprandial mortality is attributed to *κ* rather than confounded with background mortality. The probability *p*_*t,n,h*_ is naturally bounded in [0, 1] and directly separates the deterrence and preprandial mortality contributions.

##### Observation weighting

Net modes of action are estimated relative to the control feeding rate, which is poorly determined on nights with few mosquitoes. Each likelihood contribution is therefore weighted by the number of mosquitoes that fed or died preprandially in the control arm on that night (the count modelled by the control negative binomial). For an observation on day *t*, the control and intervention weights are

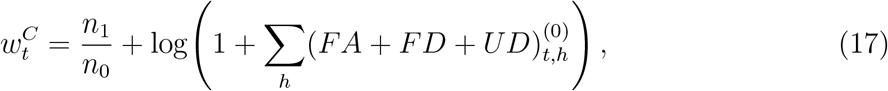

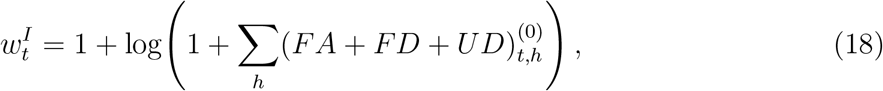

where *n*_0_ and *n*_1_ are the numbers of observations in the control and intervention arms, respectively. Each weight is applied to both the negative binomial and binomial contributions (Equations 11, 12, 14 and 15) for that observation. Weighting by the control count rather than each arm’s own catch ties estimates to nights on which the baseline contact rate is well characterised, since intervention effects are estimated relative to control feeding. The *n*_1_*/n*_0_ factor corrects for the imbalance in the number of observations between the two arms; the addition of 1 inside the logarithm ensures that nights on which no mosquitoes fed or died preprandially in the control arm still contribute to the likelihood; and the logarithm prevents nights with exceptionally high counts from dominating estimation.

#### 2.2.3 Inference

All priors were specified to be weakly informative and consistent across parameters of comparable scale. Log-scale day-specific parameters *b*_*t*_ and *m*_*t*_ were given independent *N*(0, 1) priors, permitting approximately three-fold day-to-day variation while regularising against extreme values. Net-level mean log-scale effects 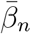 and 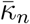 were given *N* (*−*1, 1) priors. Within-net day-to-day standard deviations 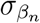 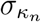 were assigned half-Cauchy(0, 1) priors. All four hut-type log-scale offsets 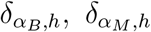, *δ*_*β,h*_, *δ*_*κ,h*_ were given (0, 1) priors, allowing up to approximately three-fold differences between hut types while remaining symmetric around no hut effect and consistent in scale across all four processes. Overdispersion parameters *ϕ*_*C*_ and *ϕ*_*n*_ were assigned half-Cauchy(0, 1) priors.

Inference was performed in Stan [15] via the `rstan` interface [16], using four Hamiltonian Monte Carlo (HMC) chains of 6,000 iterations each (3,000 warm-up). The convergence of chains was checked using the diagnostics available within Stan.

### 2.3 Postprandial mortality

Postprandial mortality is modelled separately, as it is conditional on successful feeding. For each observation, the outcome is the number of fed mosquitoes that died within 24 hours of collection (*FD*) out of the total fed (*FA* + *FD*).

#### 2.3.1 Model development

Let 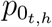 denote the baseline probability of postprandial death on day *t* in hut *h*. We model 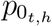 using the complementary log-log (cloglog) link,

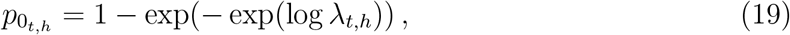

which ensures 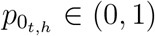 strictly and means that additive effects on the log-hazard scale, such as hut type and intervention effects, correspond to multiplicative effects on the underlying mortality rate. The log-hazard is decomposed as

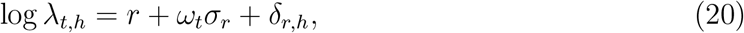

where *r* is the population-level log-hazard of baseline postprandial mortality, *ω*_*t*_ ∼**N** (0, 1) is a standardised day-specific random effect with scale *σ*_*r*_, and *δ*_*r,h*_ is a log-scale hut-type offset with the reference hut assigned zero. The quantity 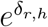 is a hazard ratio, directly interpretable as the factor by which hut *h* increases or decreases baseline postprandial mortality relative to the reference hut, independently of the day-specific baseline.

Net-induced postprandial mortality is parameterised as 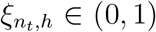, the additional probability of death among mosquitoes that survived background postprandial mortality. Since net modes of action are a property of the net itself, day-to-day variation in 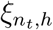 is expected to be modest relative to environmental variation in mosquito activity. Net-induced postprandial mortality is therefore modelled hierarchically, pooling information across days within each net type,

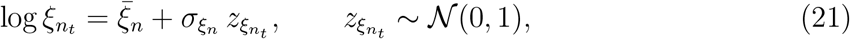

where 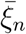 is the net-type mean log-hazard of net-induced postprandial mortality and 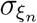 is the within-net day-to-day standard deviation. Hut-type differences in sensitivity to net-induced postprandial mortality are captured by a log-scale offset *δ*_*ξ,h*_, with the reference hut assigned zero. The hut- and day-adjusted net-induced postprandial mortality probability is then

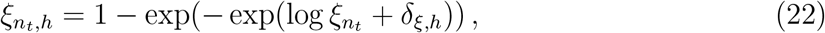

so that 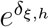 is a hazard ratio, directly interpretable as the factor by which hut *h* amplifies or attenuates net-induced postprandial mortality relative to the reference hut. The overall probability of postprandial death in the intervention arm is therefore

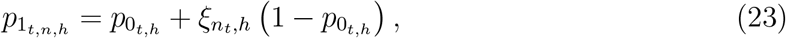

where 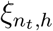 acts on the proportion of mosquitoes that survived background postprandial mortality.

#### 2.3.2 Likelihood

The likelihoods for the control and intervention arms are

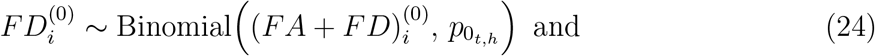

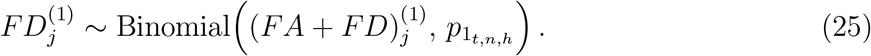

Observations are included only when at least one fed mosquito was recovered. Likelihood contributions are weighted as in the host-seeking model (Section 2.2.2), with the fed catch (*FA* + *FD*) in the control arm in place of the count of mosquitoes feeding or dying preprandially, since the postprandial likelihood conditions on having fed:

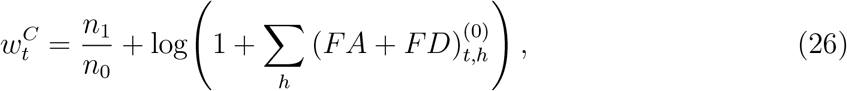

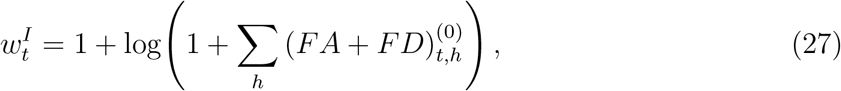

where *n*_0_ and *n*_1_ are the numbers of observations in the control and intervention arms, respectively. Weighting by the control fed catch ensures that estimates are driven by nights on which postprandial mortality can be reliably characterised from the baseline.

### 2.3.3 Inference

The population-level baseline log-hazard was given a *N* (− 3, 1) prior. The day-to-day scale parameter was given a half-Cauchy(0, 1) prior, and day-specific random effects *ω*_*t*_ ∼**N** (0, 1). The net-type mean log-hazards 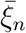 were given *N* (− 2, 1) priors, and within-net standard deviations 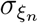 were assigned half-Cauchy(0, 1) priors. Hut-type offsets *δ*_*r,h*_ and *δ*_*ξ,h*_ were given *N* (0, 0.5) priors. Prior predictive checks confirmed that the implied distributions of 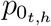 and 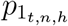 were consistent with biologically plausible values.

Similarly to the host-seeking model, inference was performed in Stan [15] via the rstan interface [16], using four Hamiltonian Monte Carlo (HMC) chains of 6,000 iterations each (3,000 warm-up). The convergence of chains was checked using the diagnostics available within Stan.

### 2.4 Vectorial capacity analysis

Following model fitting, we calculated the mean expected reduction in relative vectorial capacity for *Plasmodium falciparum* by *An. arabiensis* using the framework described in Fairbanks et al. [12]. The relative vectorial capacity is defined as the total number of potentially infectious bites that would arise from a single vector biting a single host on a single day. This was computed considering reduction in biting, preprandial mortality and post-prandial mortality. Since *β* reflects the net’s combined effect on the probability of successful feeding, of which preprandial mortality is a contributing cause, we use *β* to parameterise the reduction in feeding directly, and do not need to make the preprandial mortality adjustment included in Fairbanks et al. [12]. The relative reduction in vectorial capacity was evaluated across a range of intervention coverage levels (0–100%) to assess population-level impact.

The vectorial capacity framework was parameterised using species-specific bionomic parameters for *An. arabiensis*, the predominant vector at the study site. The probability of transmission from host to vector was 0.045 [17] and the probability of transmission from vector to host was 0.635 [18]. The human blood index was 0.47, calculated as the weighted mean of data points gathered in a systematic review [19]. The proportion of mosquitoes feeding indoors was 0.658, used as the usage parameter representing the proportion of vector bites occurring whilst the host is protected by the intervention, and was parameterised along with the proportion of parous mosquitoes in the population, 0.538, using estimates from the *AnophelesModel* R package [20]. We assume each mosquito only bites once per feeding cycle, an egg development duration of 3 days and an extrinsic incubation period of 11 days. The mortality rate is then parameterised as

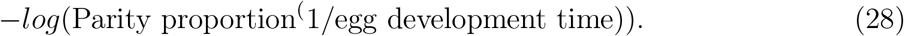

## 3 Results

### 3.1 Model fit and convergence

Both the host-seeking and postprandial models converged. The convergence of chains was checked using the diagnostics available within Stan, including the R-statistic and effective sample size [21]. Posterior predictive checks showed close agreement between observed and predicted counts for the total fed and unfed dead outcomes for the host-seeking model and fed alive outcomes for the postprandial mortality model (Figure S1), with predictions tracking the line of identity across all four hut types and in both the control and intervention arms.

### 3.2 Hut effects on baseline (control) mosquito behaviour

The median baseline feeding and preprandial mortality rates in Rapley control huts were 1.94 and 0.44, respectively (Figure S2a). Hut design had a pronounced effect on baseline feeding and preprandial mortality (Figure 1a–b). Relative to the Rapley reference hut, the East African and West African huts had substantially lower baseline feeding rates (median multiplicative effects of 0.15 and 0.05 respectively), whereas the Ifakara hut was closer to Rapley (median multiplicative effect 0.58). In contrast, the baseline postprandial mortality probability was largely insensitive to hut design, with the credible intervals for all three huts overlapping one (Figure 1c). Postprandial mortality was rare in Rapley control huts, with median baseline probability 0.001 (Figure S2b).

**Figure 1:**
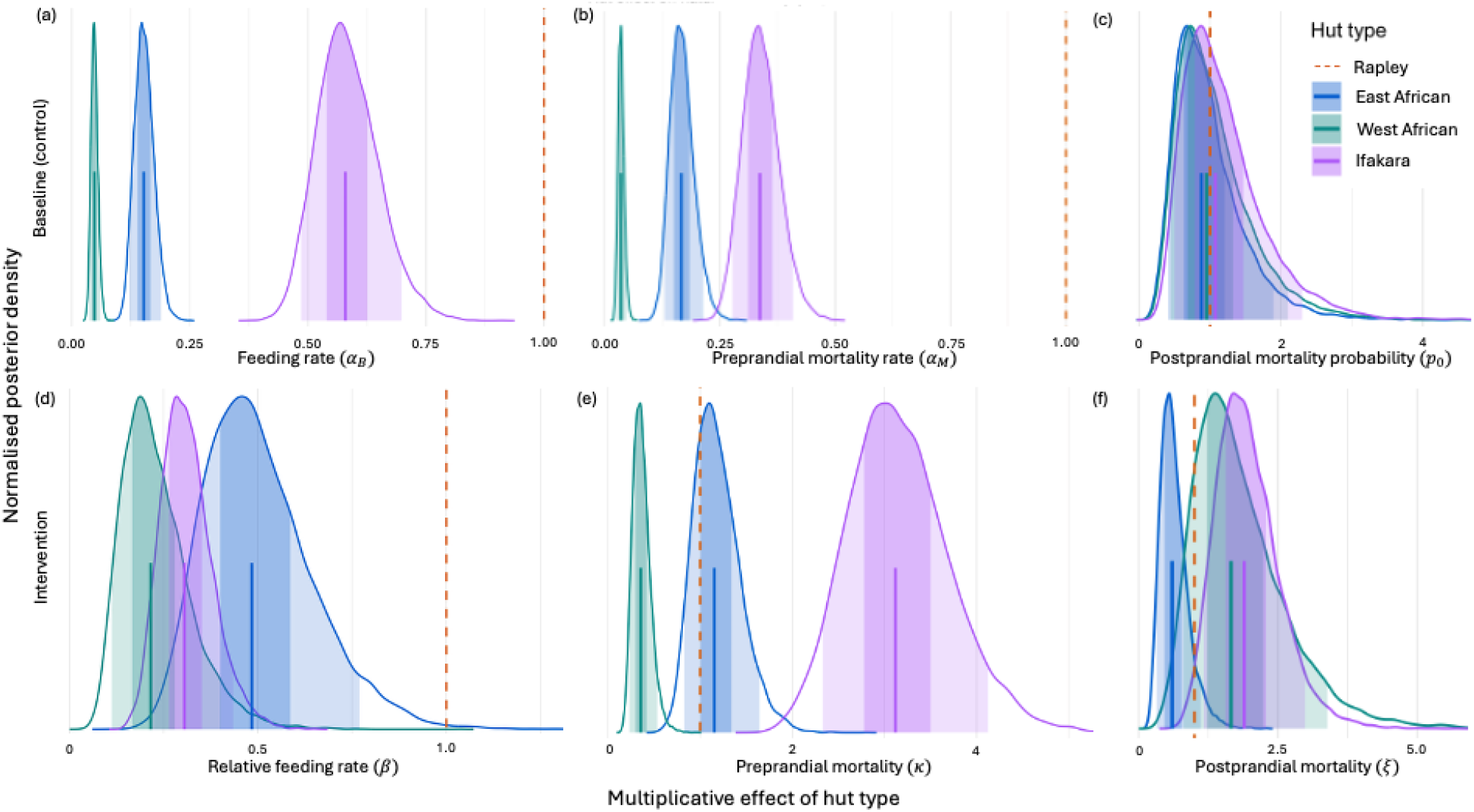
Posterior distributions of the multiplicative effect of hut type on baseline (control) (a) feeding and (b) preprandial mortality rates, (c) baseline postprandial mortality probability, and intervention (d) relative feeding rate, (e) preprandial mortality and (f) postprandial mortality, relative to the Rapley reference hut. Shaded bands denote the 50% and 90% credible interval and the vertical line the posterior median.

### 3.3 Hut effects on intervention modes of action

Hut design also influenced all three intervention modes of action (Figure 1d–f). The multiplicative hut effect on the relative feeding rate was below one for all three huts, indicating a more apparent reduction in feeding than in the Rapley hut, and was lowest in the West African hut (Figure 1d). For preprandial mortality, the Ifakara hut amplified the effect relative to Rapley (median multiplicative effect 3.12), while the West African hut reduced the effect (Figure 1e). The hut effect on postprandial mortality was more modest, with the credible intervals overlapping one for most huts and the Ifakara hut showing the largest central estimate (Figure 1f).

Net-induced postprandial mortality estimates for the West African hut were comparatively imprecise, reflecting the small number of fed mosquitoes recovered in the intervention arm; for this hut the estimates are informed largely by the hierarchical prior rather than by direct observation and should be interpreted with caution.

The net-specific magnitudes of these effects are shown in Figure S3 (relative feeding rate), Figure S4 (preprandial mortality) and Figure S5 (postprandial mortality).

### 3.4 Predicted reductions in vectorial capacity

The hut-level differences in entomological impact were translated into the predicted reductions in relative vectorial capacity (Figure 2). For every net, the predicted reduction at a given coverage was greatest in the Ifakara hut, followed by the East African hut, and smallest in either the the West African hut or Rapley huts. The magnitude of the between-hut difference was substantial: at 50% coverage, the median predicted relative vectorial capacity remaining ranged from 17.93% in the Ifakara hut to 41.10% in the West African hut for new PRONet^®^ Duo. These results indicate that the choice of experimental hut design can materially alter the predicted population-level impact of an otherwise identical intervention.

**Figure 2:**
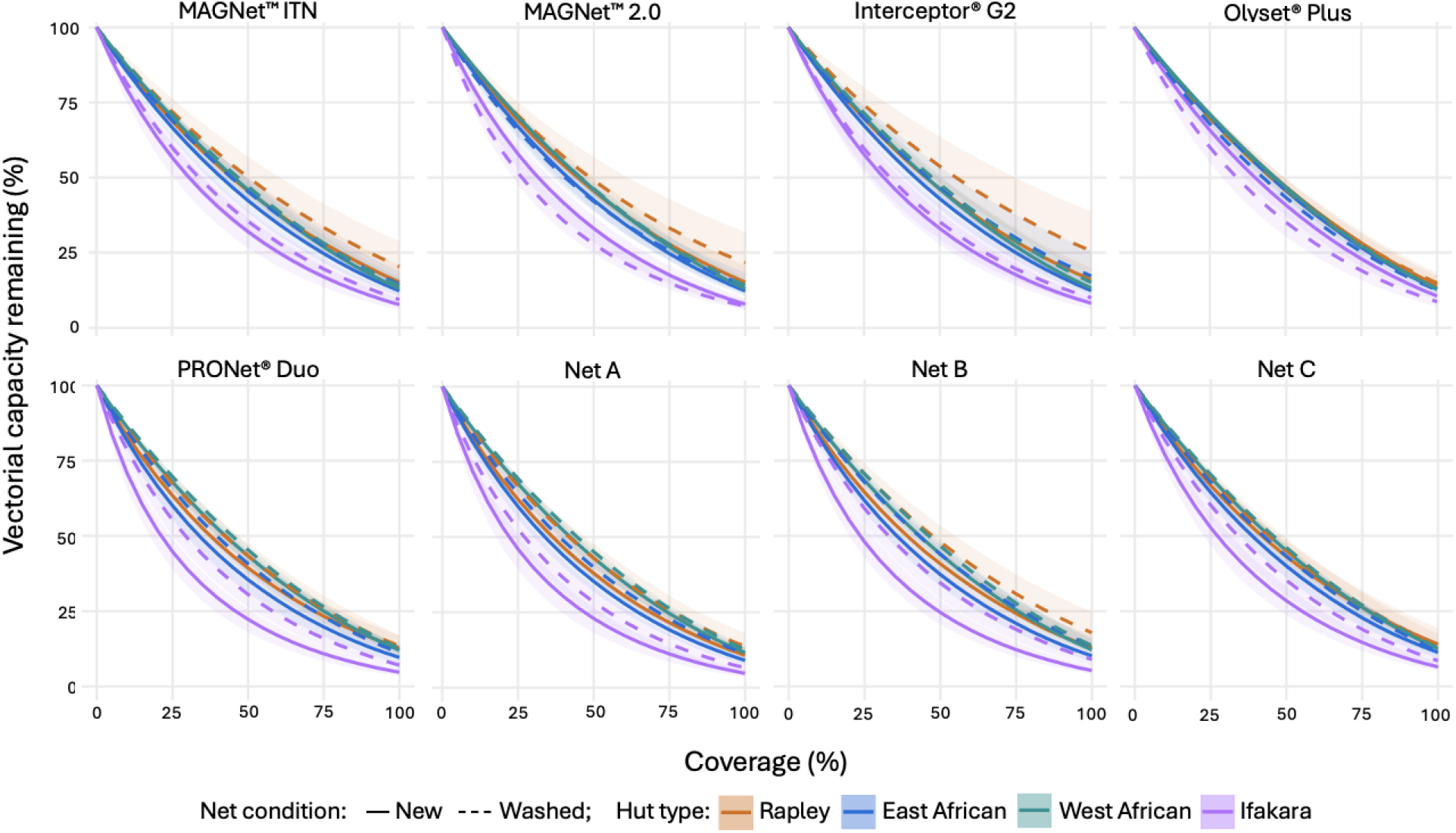
Predicted reduction in vectorial capacity across varying intervention coverage (0– 100%) for each net evaluated. Lines show the posterior median prediction for each hut type, with new nets drawn as solid lines and nets washed-20-times as dashed lines, and shaded bands denoting the 50% and 95% prediction intervals.

The contrast between new and washed-20-times conditions was generally smaller than the contrast between hut types, and its direction varied between nets: for MAGNet™, Interceptor^®^ G2, PRONet^®^ Duo and anonymised Net A,, Net B and Net C, washing reduced the predicted impact, whereas for Olyset^®^ Plus washed nets had more predicted impact, and for for MAGNet™ 2.0 results varied between hut type.

## 4 Discussion

In this study we developed a Bayesian hierarchical framework to quantify how experimental hut design shapes the entomological evaluation of ITNs. The framework jointly estimates the three modes of action through which ITNs reduce vectorial capacity: biting deterrence, preprandial mortality and postprandial mortality. The approach yields net-specific estimates for impact on modes of action and translates them into predicted reductions in vectorial capacity. We demonstrated that experimental hut design exerts an influence on every mode of action, and consequently on the predicted population-level impact.

The present analysis considers the three modes of action most commonly measured in experimental hut trials: reduction in feeding, preprandial mortality and postprandial mortality. However, the newer generation of dual active ingredient nets incorporates additional mechanisms, including sterilisation effects, delayed mortality and enhanced spatial repellency [22– 24]. The vectorial capacity framework presented here could be extended to accommodate these mechanisms through the addition of parameters capturing effects on fecundity, oviposition behaviour and mortality operating over longer timescales, offering a flexible framework for the evaluation of novel vector control interventions as they progress through the evaluation pipeline.

Mortality here is assessed at 24 hours, the window most relevant to biting on the following night and consistent across arms. While the majority of chlorfenapyr-induced mortality occurs within this window, some accrues over the subsequent days [25], and assessment at 72 hours is the established convention for chlorfenapyr nets [26]. The 24-hour window may therefore under-represent the full killing effect of slow-acting active ingredients such as chlorfenapyr. Future work could examine how the estimated modes action across choices of mortality assessment time affects resulting predictions in reduction of vectorial capacity.

The hierarchical structure of the model ensures that data-sparse huts borrow strength from the overall posterior in a principled way [27]. This is particularly relevant for the West African hut, in which very few fed mosquitoes were recovered in the intervention arm across almost all net types Figure 1. As a consequence, net-specific postprandial mortality estimates for the West African hut are largely informed by the hierarchical prior on average effects of the net considered and the hut offset rather than by direct observation. The posterior for postprandial mortality effects of nets in the West Africa hut should therefore be interpreted with caution.

Assenga et al. [3] considered two entomological endpoints, feeding and mortality (combining preprandial and postprandial) and modelled these using mixed-effects regression. The patterns are consistent with this analysis. Both analyses found higher baseline feeding rates estimated for the Rapley and Ifakara huts, relative to the East African and West African designs, and the relative ranking of products was preserved across hut designs. The mechanistic decomposition in this study provides further interpretation. Preprandial mortality acts before feeding and therefore has a greater effect on transmission than postprandial mortality. In this analysis, the West African hut produced the smallest and the Ifakara hut the largest predicted reduction in vectorial capacity, despite the high mortality measured in the West African huts. This indicates that aggregate endpoints such as total mortality may not reflect population-level impact, and supports decomposing entomological mortality outcomes into separate preprandial and postprandial modes of action.

The postprandial mortality model uses a complementary log-log (cloglog) link rather than the logit link used in Denz et al. [28] and other adaptations of this model [5, 29]. Whilst the logit link produces well-calibrated estimates of postprandial mortality probabilities, additive offsets on the logit scale do not correspond to a simple multiplicative effect on the probability scale. The implied ratio 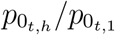 depends on the baseline probability 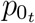, making hut effects difficult to interpret independently of the day-specific baseline. By contrast, the cloglog link models mortality as arising from a latent hazard process [30, 31], and additive offsets on the log-hazard scale correspond exactly to multiplicative effects on the hazard, i.e hazard ratios, which are interpretable independently of the baseline. This is the same principle underlying the Cox proportional hazards model in survival analysis [30], and produces hut effect estimates 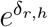 and 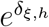 that are directly comparable across nets and days without reference to a specific baseline value. Prior predictive checks confirmed that the cloglog link produced biologically plausible distributions for both baseline and intervention-induced postprandial mortality across all hut types.

A further limitation concerns the use of fixed bionomic parameters used to translate entomological effects into reductions in vectorial capacity. Quantities such as the human blood index, proportion of bites taken indoors and parous proportion vary between vector populations and settings. The framework can be applied to a specific setting using local bionomics to yield predictions tailored to the local vector population.

Relatedly, validation of the platform comparison itself is constrained by the available data. Validation would be strengthened by data from multiple sites at which more than one hut design was deployed for the same intervention, enabling assessment of whether the relationship between platforms is consistent across vector populations and settings, rather than confounded with site-specific effects.

By quantifying the influence of hut design, this framework offers a route to more comparable and policy-relevant evidence from experimental hut trials. Because it expresses modes of action as separate, mechanism-specific parameters, its estimates can be carried directly into the mathematical transmission models used to project epidemiological impact. The substantial effect of hut design on predicted impact suggests that, when entomological trial data are used to forecast the population-level effect of a tool, the hut design under which those data were collected should be taken into account. As dual active ingredient nets and other novel interventions move through the evaluation pipeline, methods that account for hut design would help to ensure that deployment decisions rest on the properties of the interventions rather than the conditions under which they have been evaluated.

## Ethical approval

This is a secondary data analysis. No ethical approval was required.

## Consent for publication

Not applicable

## Data availability and materials

Materials will be made available on GitHub with a version DOI from Zenodo on publication.

## Author contributions ELF

Methodology, Software, Validation, Formal analysis, Investigation, Resources, Data Curation, Writing - Original Draft, Writing - Review & Editing, Visualization, Project administration. **AA:** Conceptualization, Resources, Data Curation, Writing - Review & Editing, Project administration. **OGO:** Conceptualization, Resources, Data Curation, Writing - Review & Editing. **RN:** Conceptualization, Writing - Review & Editing. **JM:** Conceptualization, Resources, Writing - Review & Editing. **SJM:** Conceptualization, Resources, Writing - Review & Editing, Project administration, Funding acquisition

## Acknowledgements

ELF acknowledges support from a University of Manchester Healthier Futures Fellowship and L’Oréal-UNESCO For Women in Science Award.

