## Supplementary figures and images for "Quantifying the impact of experimental hut design on intervention evaluation outcomes and predicted reductions in vectorial capacity"

### Figure S1

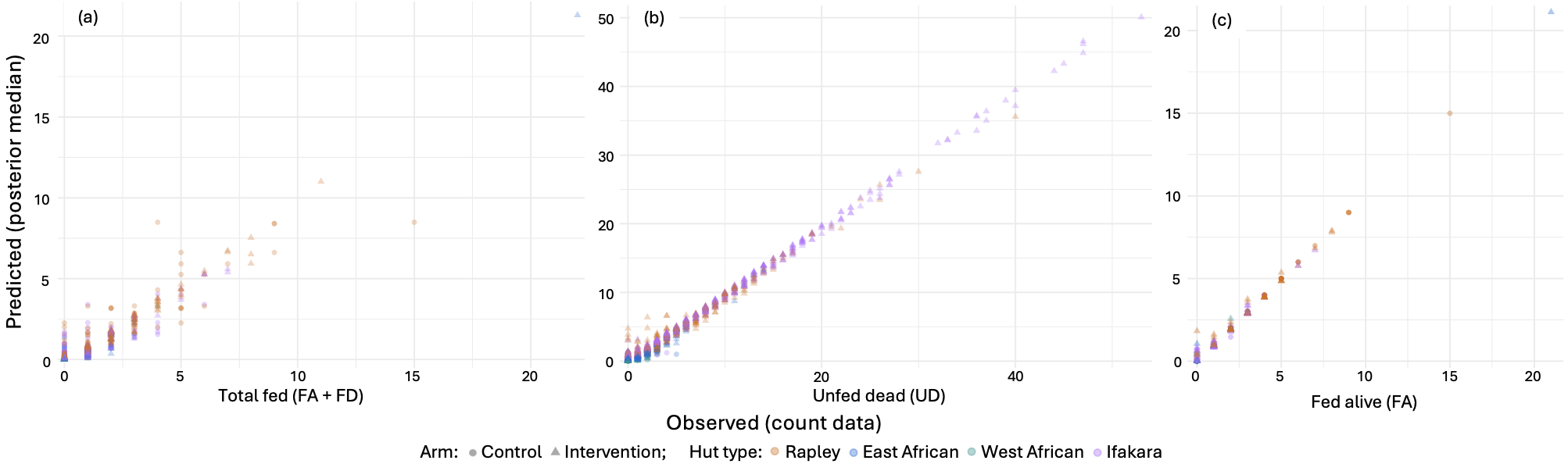

### Figure S2

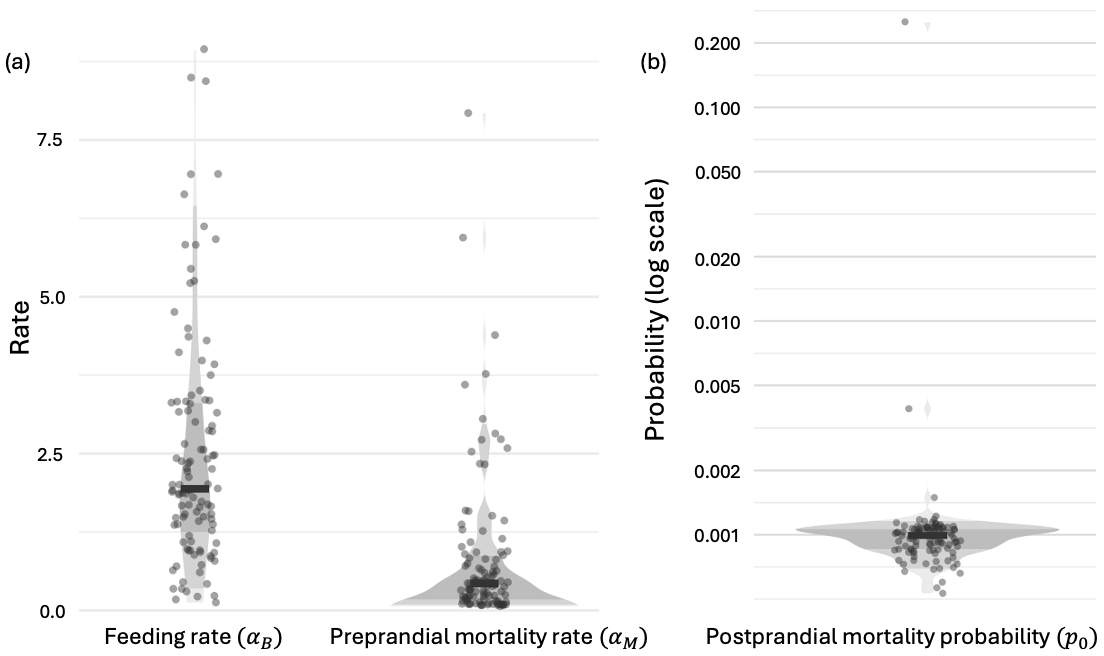

### Figure S3

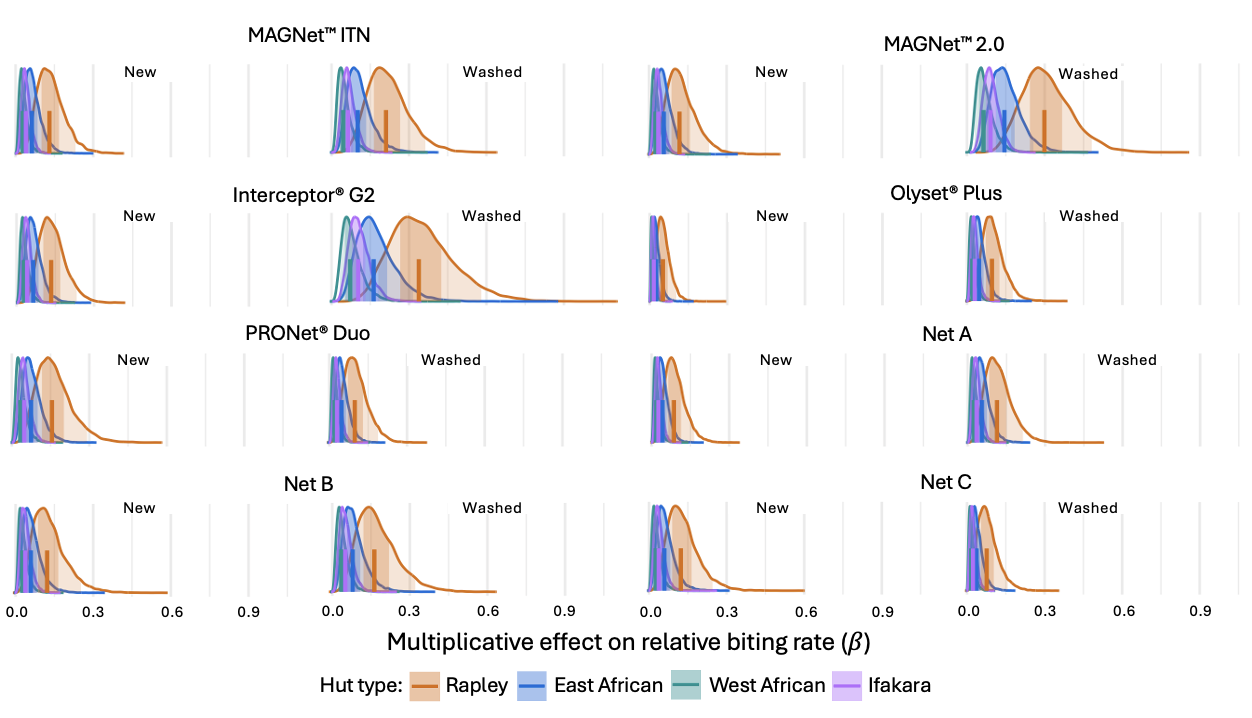

### Figure S4

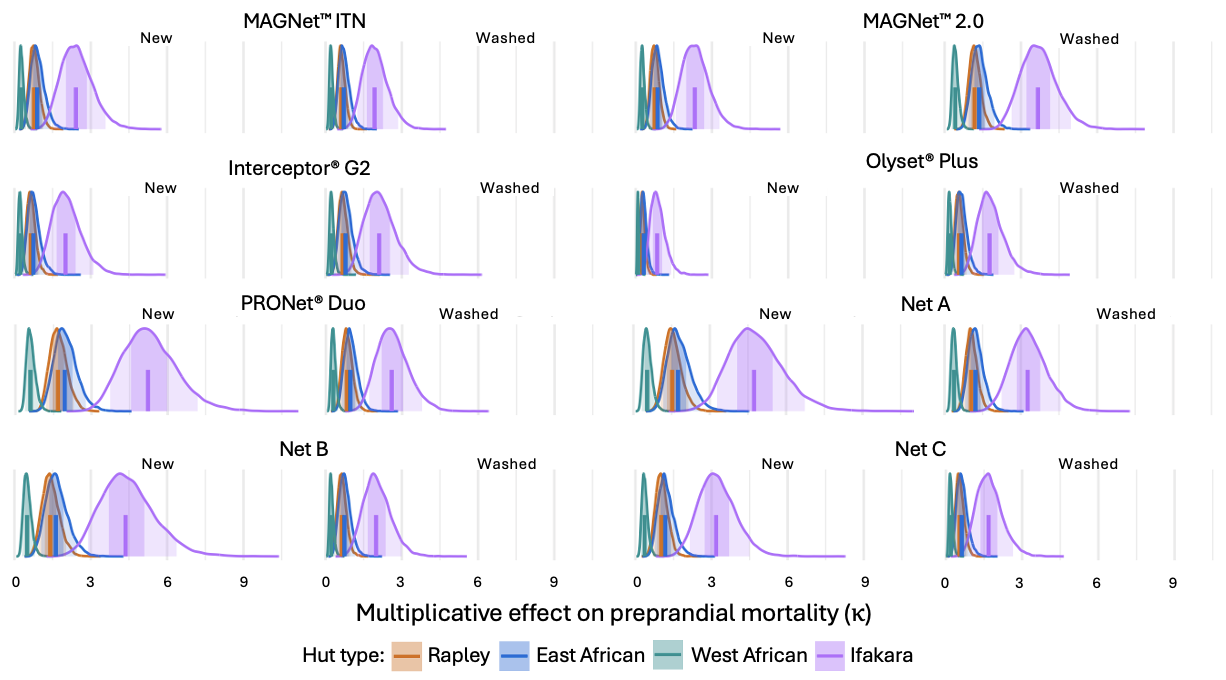

### Figure S5

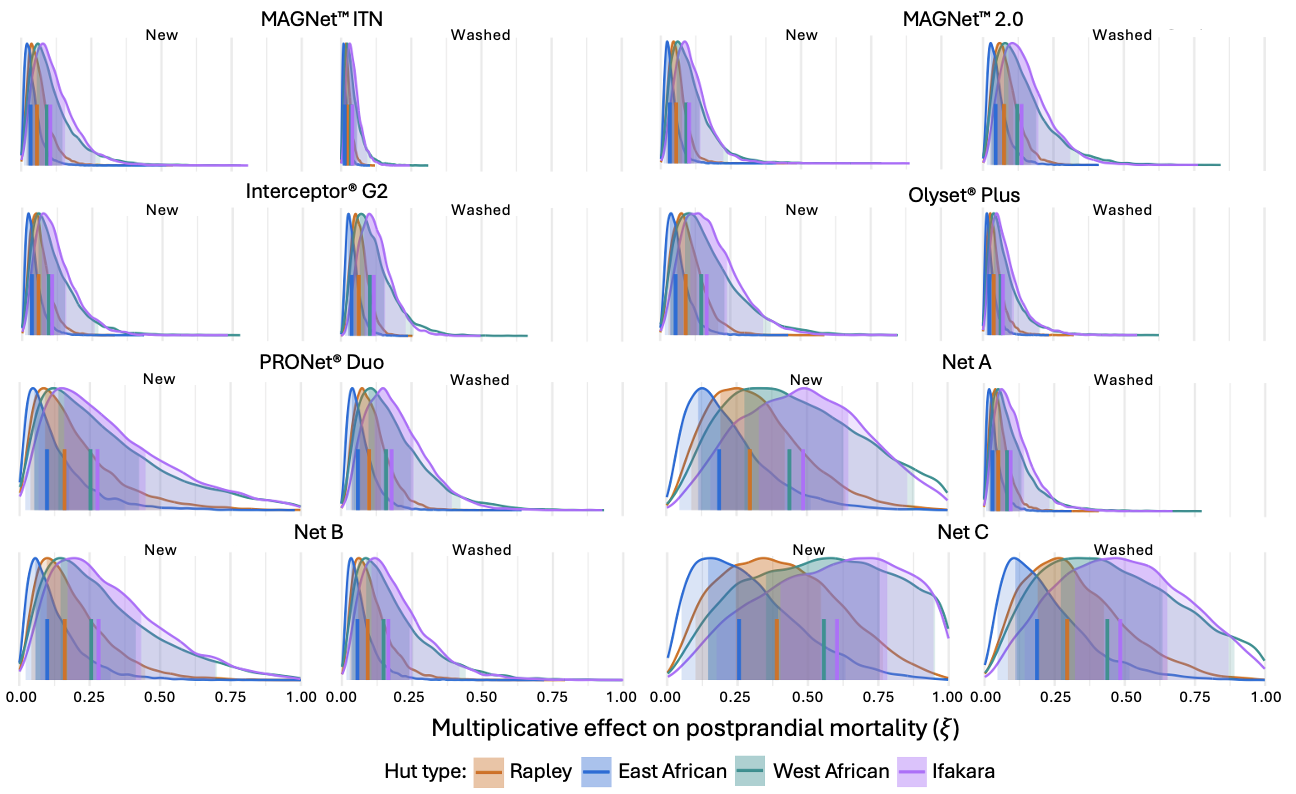
